# Cortical neural activity during responses to mechanical perturbation: Effects of hand preference and hand used

**DOI:** 10.1101/2024.11.26.625431

**Authors:** Kevin Hooks, Kimia Kiani, Qiushi Fu

## Abstract

Handedness, as measured by self-reported hand preference, is an important feature of human behavioral lateralization that has often been associated with hemispheric specialization. We examined the extent to which hand preference and whether the dominant hand is used or not influence the motor and neural response during voluntary unimanual corrective actions. The experimental task involved controlling a robotic manipulandum to move a cursor from a center start point to a target presented above or below the start. In some trials, a mechanical perturbation of the hand was randomly applied by the robot either consistent or against the target direction, while electroencephalography (EEG) was recorded. Twelve left-handers and ten right-handers completed the experiment. Left-handed individuals had a greater negative peak in the frontal event-related potential (ERP) than right-handed participants during the initial response phase (N150) than right-handed individuals. Furthermore, left-handed individuals showed more symmetrical ERP distributions between two hemispheres than right-handed individuals in the frontal and parietal regions during the late voluntary response phase (P390). To the best of our knowledge, this is the first evidence that demonstrates the differences in the cortical control of voluntary corrective actions between left-handers and right-handers.

## 1. INTRODUCTION

Handedness is a well-known form of behavioral lateralization (or asymmetry), which can be measured as differences in hand preference or hand performance between two limbs (Nicholls et al., 2010; Scharoun and Bryden, 2014). Between these two, self-reported hand preference is more commonly used in handedness research due to its simplicity in assessment. For example, the well-established Edinburgh Handedness Inventory (EHI) uses a variety of tasks, such as writing, sport, and manual labor to determine which hand an individual would perform those tasks with. A score, the Laterality Quotient, is then given to quantify the degree of handedness of the individual (Oldfield, 1971). It has been reported that approximately 10% of the population is left-handed (Papadatou-Pastou et al., 2020), measured as hand preference.

Moreover, the frequency of handedness can vary between different demographics since gender, age, and nationality have all been shown to play a role (Gilbert and Wysocki, 1992; Ida and Bryden, 1996; Faurie et al., 1999).

The complete explanation for the prevalence of right-handedness remains unclear (in the present work we use handedness as defined by self-reported direction of hand preference). Nevertheless, extensive neuroimaging research has demonstrated that handedness is associated with various structural differences in the cerebral cortices of adults (Toga and Thompson, 2003; Hervé et al., 2006; Luders et al., 2010; Andersen and Siebner, 2018; Howells et al., 2018; Sha et al., 2021; Chormai et al., 2022). Furthermore, it is often found that right-handed individuals have stronger asymmetrical activation patterns across two hemispheres, i.e., neural lateralization, than left-handed individuals. For instance, functional neuroimaging has revealed that bilateral activation patterns are more likely to occur in left-handers during cognitive functions, such as language-processing tasks (Pujol et al., 1999; Knecht et al., 2000), face-processing tasks (Frässle et al., 2016; Rossion and Lochy, 2022), and body-perception tasks (Karlsson et al., 2022). For motor tasks, left-handers were found to be associated with more symmetrical fMRI activation patterns than right-handers during finger-tapping (Kim et al., 1993; Tzourio Mazoyer et al., 2015), sequential finger movement (Solodkin et al., 2001), repetitive ball squeezing (Crotti et al., 2022), and pantomiming familiar object manipulations (Vingerhoets et al., 2012). Connectivity between cortical areas was also found to be more symmetrical in left-handers than right-handers during fist closure movements (Pool et al., 2014).

Electroencephalography (EEG) has also been used as an imaging tool to understand the neural substrates of handedness in motor tasks. The readiness potential before simple movement was found to be more lateralized in right-handers than in left-handers (Schmitz et al., 2019). EEG connectivity in the parietal-premotor networks was more bilateral in left-handers than in right-handers during action observations. Moreover, motor imagery tasks were found to be associated with less lateralized alpha band (8-13 Hz) power in left-handers than right-handers (Zapała et al., 2020), but opposite results have also been reported (Lajtos et al., 2023). Additionally, alpha band power was found higher in the left frontal area of left-handers but higher in the right frontal area of right-handers during the performance of motor tasks from the EHI assessment (Packheiser et al., 2020). However, most existing EEG-based research on the handedness effect in motor control has focused on well-practiced or predictable motor tasks. In contrast, little is known about the extent to which handedness (or hand preference) affects neural lateralization in the sensorimotor system during unpredictable situations where motor actions may need to be altered in response to perturbations. The ability to use sensory feedback to correct deviations that interfere with task goals is critical to performing daily activities as unforeseen events may occur, e.g., someone accidentally bumps your arm when you are holding a cup of drink. Therefore, it is important to advance our understanding of the role of handedness in feedback-driven corrective voluntary control.

Mechanical perturbation is a useful probe for corrective control (Scott, 2012). In this study, we used EEG in a classic experimental paradigm to investigate the neural response underlying corrective actions during an unimanual motor task with mechanical perturbations. Human participants held a robotic device with the goal of quickly moving to either an upward or a downward target. A mechanical perturbation would occur in the same or opposite direction of the target during random trials, requiring corrective responses after the perturbation. Previous studies using electromyography (EMG) revealed that the motor response to such unexpected environmental changes consists of a series of three components after the onset of the perturbation (Pruszynski et al., 2011; Forgaard et al., 2016): a short-latency reflex (< 50 ms), a long-latency response (50 – 120 ms), and voluntary corrections (> 120 ms). Importantly, both human and non-human primate studies have demonstrated that the cerebral cortex is involved in the goal-dependent modulation of the second and third components (Shemmell et al., 2009; Spieser et al., 2010; Pruszynski et al., 2014). EEG is particularly suitable for investigating cortical activities in the short duration of corrective control given its high temporal resolution.

However, existing EEG studies that used similar mechanical perturbation paradigms only examined the dominant hand of right-handed individuals with a focus on the second response component, i.e., < 100 ms post-perturbation (Mackinnon et al., 2000; Kourtis et al., 2008). A more recent study showed that the late (100-350 ms) cortical response to mechanical perturbation of the right wrist of right-handers activates a larger network than the early (20-100 ms) response (Yang et al., 2017). Consistent with earlier findings, they revealed that the early response is centered around the contralateral primary motor cortex. Furthermore, significant contributions from frontal and parietal areas were found in the late response. The present study extended the classic mechanical perturbation paradigm to examine both left and right hands in both left-handers and right-handers. Additionally, we analyzed the spatiotemporal distribution of the EEG signals over the entire 500 ms duration of the task, emphasizing the overlooked late voluntary response period. Such analyses allowed us, for the first time, to systematically determine how handedness may affect the neural activities within a broad sensorimotor network underlying the corrective motor control processes. Based on the literature on the handedness-related differences in neural lateralization, we hypothesized that right-handers, compared to left-handers, would be associated with more asymmetrical spatial patterns of event-related potentials (ERPs) during the voluntary control of corrective responses.

## 2. MATERIALS & METHODS

### 2.1 Participants

A total of 22 participants completed the experiment. They had normal or corrected-to-normal vision and no history of musculoskeletal or neurological disorders. All participants were naïve to the purpose of the study and gave informed consent to participate in the experiment. The experimental protocols were approved by the Institutional Review Board at the University of Central Florida in accordance with the Declaration of Helsinki. The handedness of the participants was evaluated with the Edinburgh Handedness Inventory (EHI), which is a questionnaire that asks about the hand preferences in 10 different common motor tasks (Oldfield, 1971). The EHI produces a score between -100 and 100, representing the directional preference difference between the right and left hands. A cutoff EHI score of 0 was used to assign the participants to two groups (see **Supplementary Materials**). The right-handed group consisted of 10 participants (6 M, 4 F, 23.9 ± 4.9 years), and the left-handed group consisted of 12 participants (6 M, 6 F, 23.3 ± 3.9 years). Note that all participants in the right-handed group scored above 40, and 10 participants in the left-handed group scored below -40. We included two participants who scored -33 and -7 in the left-hand group, although they can also be considered ambidextrous.

### 2.2 Experimental setup

Participants were instructed to arrive with hair clean and free of styling products, i.e., mousse, hair spray, hair gel, etc. The forehead, scalp, and hair were cleaned with alcohol wipes to remove excess skin oils before the cap was placed on the participant. A 32-channel EEG ActiCap system (Brain Products, Germany) was individually fitted to each participant’s head size to ensure a proper connection of the electrodes, following the international 10-20 setup.

Conductive gel was applied to each electrode to achieve an impedance level below 20 kOhms. The EEG signals were recorded at 500 Hz.

Participants were seated comfortably in front of a table where a haptic robot Phantom Premium 1.5 (3D systems, Rock Hill, SC) and a computer monitor were located. The chair was adjusted for each individual to allow the participants to perform the experimental task with their elbows resting on the armrest of the chair with approximately 90 degrees of flexion (**Fig 1A**).

**Figure 1.**
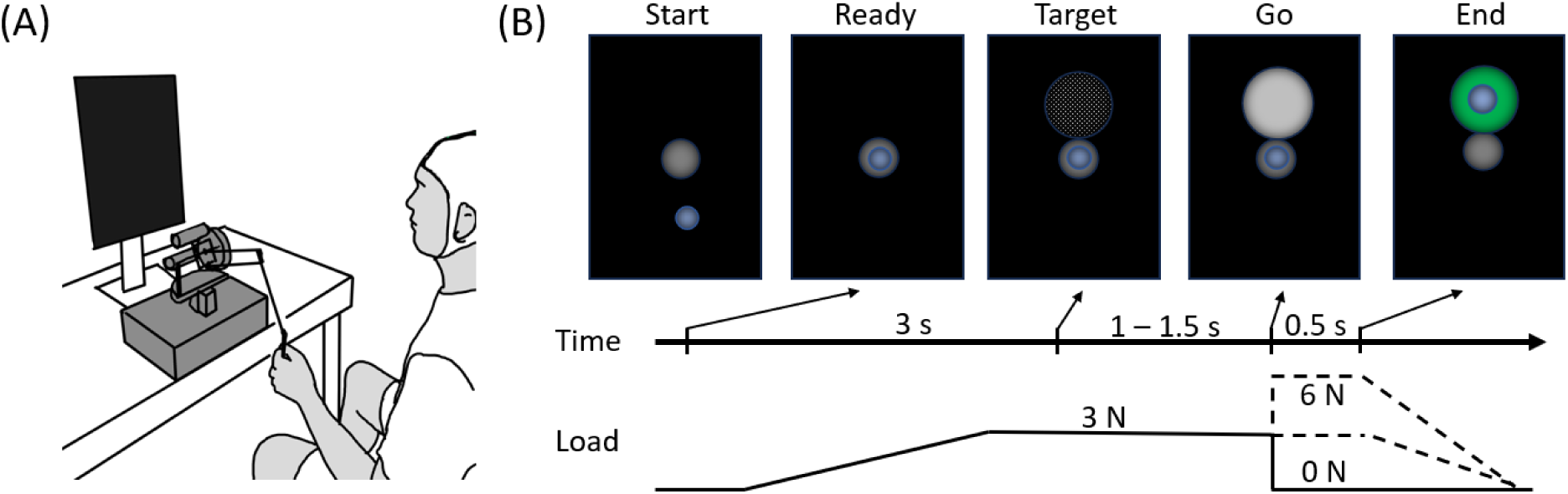
(A) Experimental setup**. (B)** Visual feedback and event timing in one trial.

Participants were instructed to hold the end-effector of the robot vertically and firmly and follow the visual feedback displayed on the monitor. The visual feedback was rendered at 60 Hz and it was positioned at eye level and at approximately 1 m distance. The vertical position of a participant’s hand was measured through the robot at 500 Hz, which was used to control the movement of a small blue sphere (2.5 cm diameter) on the display screen that will be referred to as the cursor. Participants were instructed to move the robot end-effector with elbow and wrist motion. The experimental task was implemented using a customized program with the CHAI3d library (Conti et al., 2005). A soft virtual spring-damper (spring constant 0.25 N/mm, damping constant 0.03 N·s/mm) constraint was rendered by the robot to limit the movement of the hand in left-right and forward-backward directions. The movement recording and EEG signals were synchronized by event triggers generated in a customized LabView program (National Instruments, Austin, TX).

### 2.3 Experimental Procedure

Participants followed a series of visual cues (**Fig 1B**). To initiate a trial, participants moved into a gray Start sphere (4 cm diameter). After staying inside the Start sphere for 0.5 s, a 3 N background load was gradually applied by the robot in the upward direction, and the participants were instructed to exert a downward force to maintain the cursor inside the Start sphere. The ramping of the background load was completed in 2 s. After another 0.5 s, a white mesh Target sphere (16 cm diameter) appeared adjacent to the Start sphere, either above or below, indicating which direction the participant must move the cursor. The Target sphere then turned solid grey after a random interval of 1 to 1.5 s, which was defined as the Go cue. At the time of the Go cue, the robot may behave in one of the following three ways: No perturbation (N), Upward perturbation (U), or Downward perturbation (D). Participants were instructed to move the cursor into the target sphere within 0.5 s, regardless of how the robot may act. For N conditions, the background load remained unchanged. For U and D conditions, the background load was quickly ramped to 6 N and lowered to 0 N in 20 ms, respectively. Therefore, there were 6 total experimental conditions as follows: Up target/Up perturbation (UU), Up target/No perturbation (UN), Up target/Down perturbation (UD), Down target/Up perturbation (DU), Down target/No perturbation (DN), and Down target/Down perturbation (DD). A trial was considered successful if the cursor remained inside the Target sphere after 0.5 s with less than 0.01 m/s velocity. The target sphere changed to green for success or red for failure of the trial, and the non-zero background load was gradually removed. The type of abrupt mechanical perturbations we used in this study was known to elicit different corrective motor responses that are dependent on the congruency between the target and perturbation directions (Pruszynski et al., 2011).

There were 4 blocks of 50 trials. The inter-trial and inter-block intervals were 3 s and 1 min, respectively. The participants alternated right and left hands for each block (order counterbalanced). Within each block, the 6 experimental conditions were randomly presented. Each block contained 5 trials of UN and DN non-perturbation conditions, and 10 trials of DD, DU, UD, and UU perturbation conditions, respectively. Therefore, there were a total of 10 trials in each non-perturbation condition and 20 trials in each perturbation condition for each hand.

### 2.4 Data Analysis

All data was processed in MATLAB (Mathworks, Natick, MA). The vertical positions of the cursor were low-pass filtered by zero-lag Butterworth 4-th order 5 Hz filter. To examine if hand preference and whether the dominant hand was used modulated the behavioral response after perturbation, we divided the [0, 500] ms time window after Go cue into twenty 25 ms segments. First, we examined the difference in motor behavior in relation to the perturbation regardless of the target by computing the perturbation-related response difference. Within each segment, the average vertical position of the trials with the Down perturbation was subtracted from that of the trials with the Up perturbation, i.e., (UU+DU)/2‒(UD+DD)/2. A mixed two-way ANOVA was implemented for each segment with Handedness being a between factor and Hand being a within factor. Second, we examined the difference in motor behavior that aimed to bring the cursor to the target despite the mechanical perturbation. This was quantified as the average vertical position of the trials with the Up target subtracted by the average vertical position of the trials with the Down target, i.e., (UU+UD)/2‒(DU+DD)/2. Mixed two-way ANOVA with Handedness being a between factor and Hand being a within factor was again implemented for each segment. Because we did not find significant effects of either Handedness or Hand in all segments, we took the target-related response differences grand average across hands and pooled all participants to perform a series of one-sample t-tests against zero to detect the onset of target-specific response. Specifically, we examined ten 10 ms segments starting between 100 and 200 ms after Go cue. This was because it was well known that target-specific kinematic response in this type of perturbation paradigm starts after 100 ms (Pruszynski et al., 2008). Bonferroni correction was used to account for multiple hypothesis testing.

EEG data was pre-processed using the EEGLAB toolbox (Delorme and Makeig, 2004). Firstly, the data was re-referenced to the common average and band-filtered between 1 - 50 Hz using a windowed type I linear phase sinc FIR filter (Widmann et al., 2015). Next, automatic subspace reconstruction (ASR) was used to remove high-amplitude artifacts from the data (Mullen et al., 2013). This method uses principal component analysis to determine if portions of the data exceed a statistical threshold for rejection and can interpolate the data based on surrounding channels. Any channels or portions of data that are rejected by the algorithm were replaced using spherical interpolation of the nearest 4 channels. Epochs were then created for [- 2, 2] s with respect to the Go cue, i.e., the onset of the perturbation. An adaptive mixture independent component analysis was used to compute the independent components (Palmer et al., 2012) which was used to reject components that are likely to be artifacts by the SASICA toolbox (Chaumon et al., 2015).

The present study focuses on the generation of corrective responses during motor execution, therefore we excluded the UN and DN conditions since perturbation was not applied in these trials (for data analysis of UN and DN trials, see **Supplementary Materials**). For the four conditions with perturbation, we pooled the trials together to obtain the average ERP of each hand for each participant. This was because previous works showed that the congruency of the target and perturbation directions did not lead to different ERP waveforms at least for the first 150 ms after perturbation onset (Mackinnon et al., 2000). Additionally, fMRI revealed that the preparation of resisting or letting go of a mechanical perturbation may involve the same sensorimotor network even though the motor outputs of these two behaviors are different (De Graaf et al., 2009).

The average ERP was first analyzed using topographic analysis of variance (TANOVA) with the Ragu toolbox (Koenig et al. 2011). TANOVA is a non-parametric randomization test that detects the topographic dissimilarity between spatial distributions of ERP, without a-priori bias in selecting regions of interest or time windows (Murray et al., 2008). Such dissimilarities provide a statistical means of determining if and when the underlying neural circuits may differ between conditions or groups. Specifically, we used a 20 ms shifting time window with a step size of 10 ms across 0 – 500ms post Go cue. All channels were included except Fp1 and Fp2. We used a two-way mixed design, with the between factor being Handedness and the within factor being the acting Hand.

Based on the TANOVA results and visual examination of the spatiotemporal patterns of the ERP, as well as the literature about cortical responses after movement perturbation, we chose to focus on two temporal regions of interest (ROI): 140-160 ms and 380-400 ms post Go cue. The goal of the follow-up ERP analysis was to compare the spatial distribution across two hemispheres. Therefore, we chose two frontal ROIs (left frontal F3, FC1, FC5; right frontal F4, FC2, FC6) and two parietal ROIs (left parietal P3, CP1, CP5; right parietal P4, CP2, CP6). The channels were grouped within each spatial ROI and the average ERP was obtained within the 20 ms temporal ROIs. We performed three-way mixed ANOVAs to investigate two within factors: Hemisphere (contralateral/ipsilateral ROI) and acting Hand (Right/Left), as well as one between factor: Handedness. Post-hoc comparisons with Bonferroni corrections were used after significant interactions were revealed.

## 3. RESULTS

### 3.1 The corrective actions after perturbation were similar across handedness groups and acting hands

We first examined the behavioral data to analyze the timing of the onset of voluntary correction in kinematics. The mechanical perturbations applied at the GO cue displace the hands in either upward or downward directions. Overall, the experimental task was easy to perform, and the mean success rate was approximately 96% across all conditions. Because of the transmission and processing time of the sensorimotor system, as well as the inertia of the musculoskeletal system, the corrective response to alter the initially programmed motor output can only be observed in the movement after a delay. When the perturbation and target directions were the same, participants had to inhibit the intended movement to avoid overshooting the target. In contrast, when the perturbation and target directions were opposite, participants had to issue a stronger motor output to the target direction to overcome the perturbation. We observed that the overall kinematic responses to the mechanical perturbation in our experiments were similar across the left and right hands, as well as across left-handed and right-handed groups (**Fig. 2A**). This was supported by the lack of significant effect of Handedness and Hand in all mixed ANOVA on the perturbation-related as well as target-related response differences. The grand averages revealed that the perturbation-related component was strong in the initial response (**Fig. 2B**), and the target-related component exerted gradually more influence in the later part of the response (**Fig. 2C**). A series of one-sample t-tests revealed that, on average, the target-related kinematic response started in the time window of 150-160 ms after the GO cue. We also performed the same kinematic analysis by excluding the two participants with -33 and -7 EHI scores, and the results remained the same. In sum, the behavioral results are consistent with previous findings about kinematic correction onset time in similar behavioral paradigms that used mechanical perturbations on moving upper limbs (Mackinnon et al., 2000; Pruszynski et al., 2008). Additionally, these results indicate that any differences in EEG signals are unlikely to be caused by differences in motor outputs or sensory inputs, but rather by differences in cortical processing.

**Figure 2.**
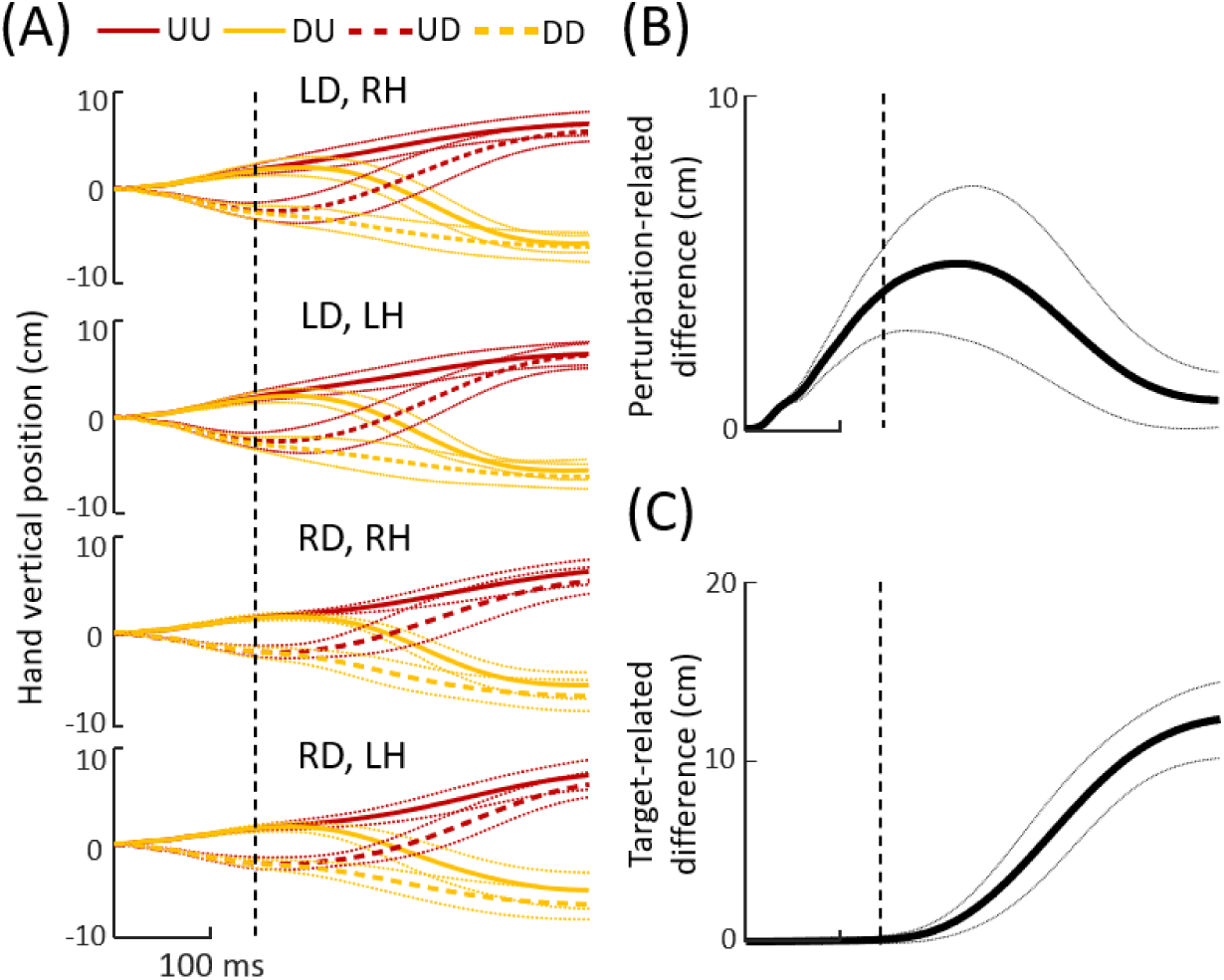
Kinematic responses after Go cue (time zero). (**A**) Vertical movement of the cursor. LD and RD denote left-handed and right-handed groups, respectively. LH and RH denote the left hand and the right hand as the acting hand, respectively. (**B**) Grand average of the difference between cursor movements with Up and Down perturbations. (**C**) Grand average of the difference between cursor movements for Up and Down targets. The thick solid and dashed lines are the means, and the thin dotted lines are the standard deviations. The vertical black dashed line represents the start of target-related kinematic responses after the initial movement driven by the mechanical perturbation.

### 3.2 Spatial distributions of ERP were different across handedness groups

We first explored the effect of *Handedness* group and acting *Hand* on the overall spatial distribution of the ERP using TANOVA (**Fig. 3A**). Significant main effects of *Handedness* were found between 110 ms and 160 ms, as well as between 380 ms and 500 ms post perturbation onset. Significant main effects of Hand were found for all 10 ms time windows after 40 ms post perturbation onset, except between 310 ms and 320 ms. A significant interaction was only found between 320 ms and 330 ms after perturbation onset. The *Hand* effect was expected because the motor control of a simple unimanual task is known to be primarily driven by the sensorimotor network contralateral to the acting hand, which leads to an asymmetrical spatial distribution of ERP between contralateral and ipsilateral hemispheres. The 40 ms early start of the cortical involvement difference between two hands (**Fig. 3B**) is consistent with the spatiotemporal distribution of the long latency response to mechanical perturbation observed in the primary motor cortex, e.g., N58-P58 (Kourtis et al., 2008). The timing of *Handedness* effects suggested that there are two periods where left-handers and right-handers differed in their cortical activities during the response to the perturbation. Visual examination of the ERP during these periods revealed that the ERP spatial pattern within each period was consistent. Therefore, we focused on two 20-ms temporal ROIs that best capture these patterns at their peaks. Specifically, the first ROI (140 – 160 ms post perturbation) was defined as the N150, which featured a strong negative potential in the frontal areas (**Fig. 3C**). The second ROI (380 – 400 ms post perturbation) was defined as the P390, which featured positive potentials in the parietal areas (**Fig. 3D**). Similar results were obtained after excluding two participants with -33 and -7 EHI scores (see **Supplementary Materials**). The following analyses within these two temporal ROIs were performed by grouping electrodes from contralateral and ipsilateral hemispheres separately (instead of left versus right hemispheres) to account for the lateralization of unimanual motor control (**Fig. 4**).

**Figure 3.**
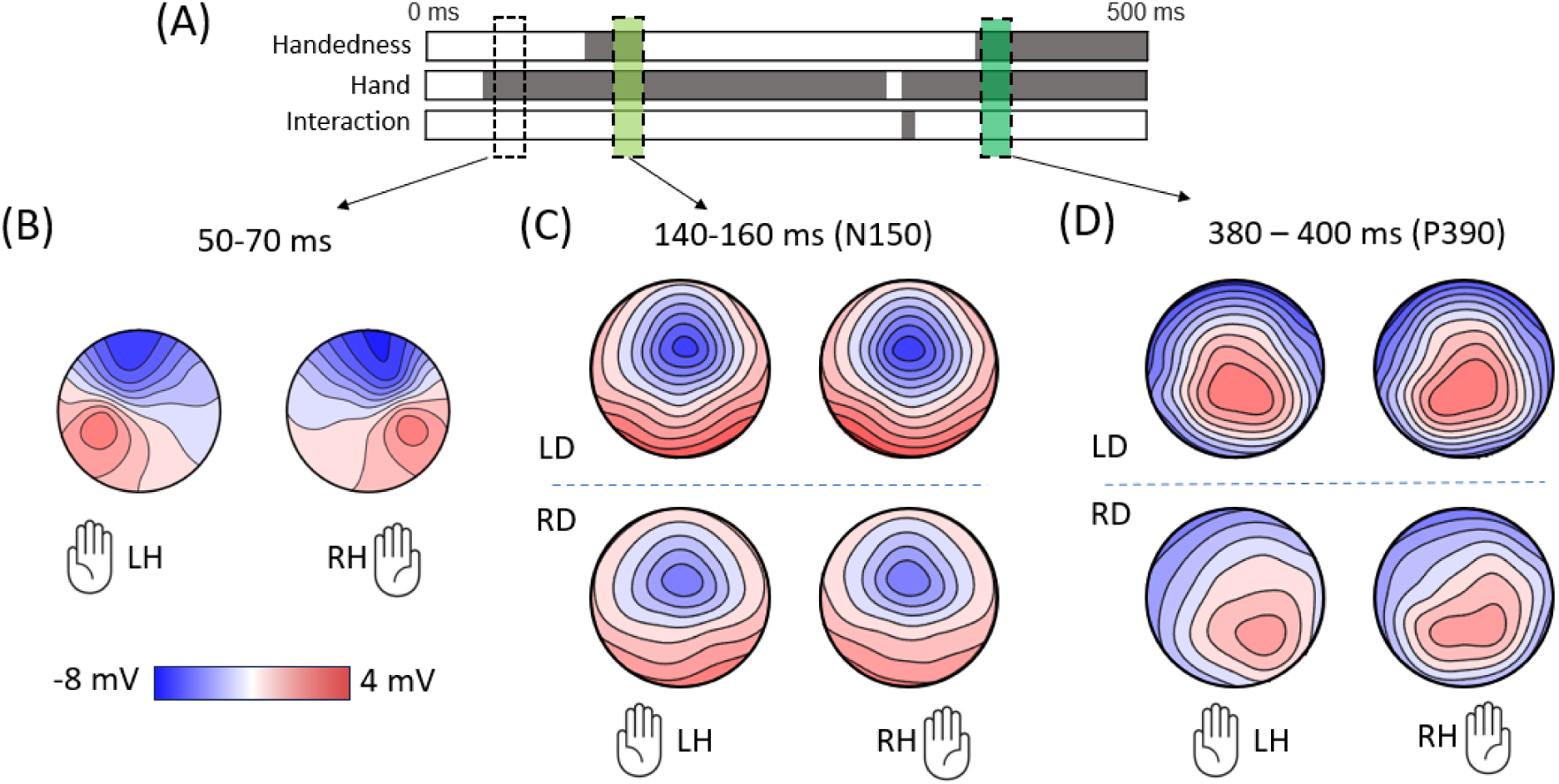
(**A**) Each horizontal bar represents results from TANOVA analysis for the 0 – 500 ms period post perturbation onset. Dark-shaded areas indicate significant effects (p < 0.05). The dashed empty box represents the 50-70 ms time window associated with the long-latency response. The light green shaded box represents the N150 temporal ROI. The darker green shaded box represents the P390 temporal ROI. (**B** - **D**) Topographic representation of the average ERP within three distinct time windows. LD and RD represent left-handed and right- handed participants, respectively. LH and RH represent the left hand and right hand being used as the acting hands, respectively.

**Figure 4.**
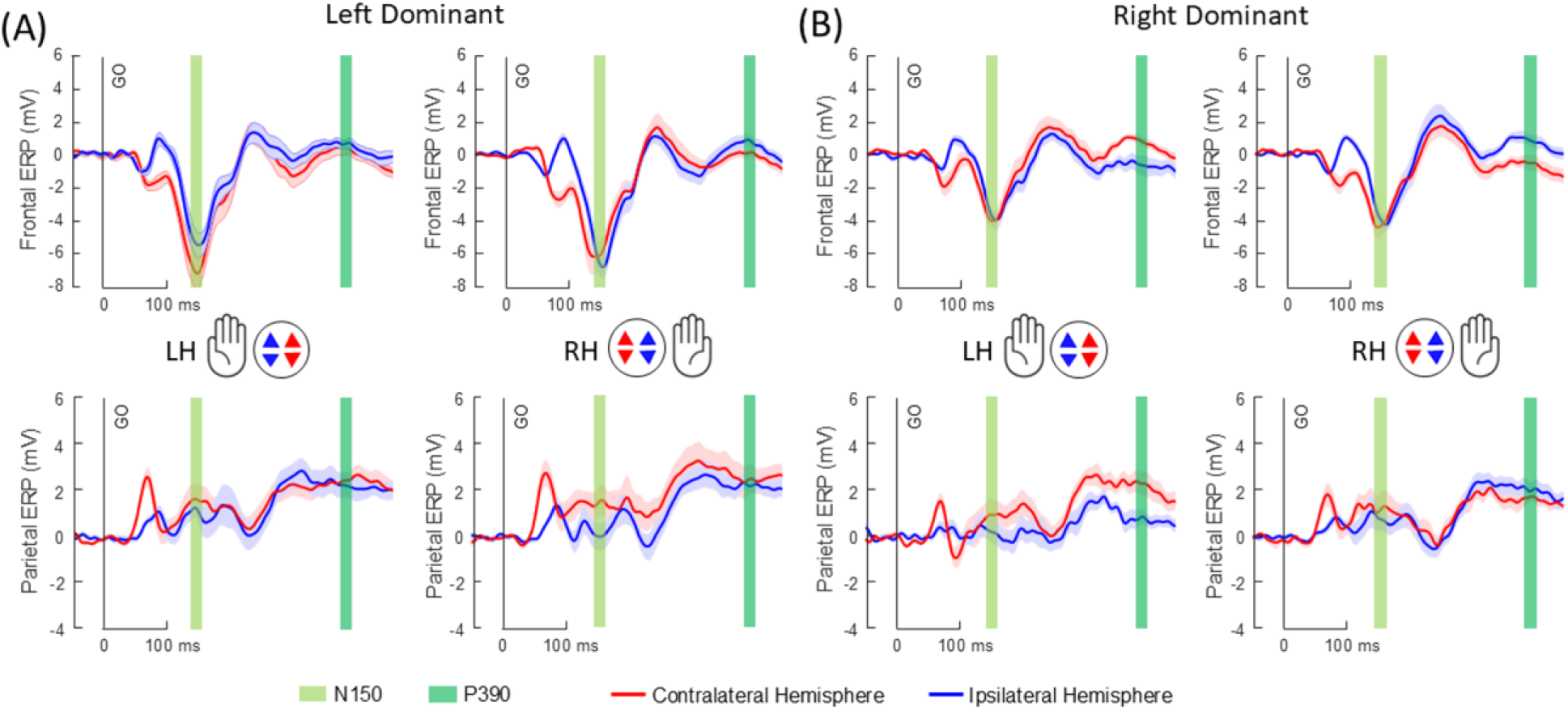
Average ERP traces from frontal ROIs (top row) and parietal ROIs (bottom row). (**A**) Left-handed participants. (**B**) Right-handed participants. Red lines and shaded areas (Mean ± S.D.) represent the contralateral hemisphere, blue lines and shaded areas represent the ipsilateral hemisphere. The first light green shaded area represents the N150 temporal ROI, and the second darker green shaded area represents the P390 temporal ROI.

### 3.3 The Left-handed group had a more negative frontal ERP than the right-handed group during N150

In the frontal ROIs, we observed that the left-handed group showed a greater negative peak in the ERP during N150 (**Fig. 4** and **Fig. 5A**). Moreover, the N150 was more negative in the contralateral hemisphere, suggesting an asymmetrical distribution of ERP with respect to the midline. This was confirmed by a three-way mixed ANOVA showing a significant effect of Handedness (F(1,20) = 6.497, p = 0.019, η_p_ ^2^ = 0.245) and Hemisphere (F(1,20) = 19.060, p < 0.001, η_p_ ^2^ = 0.488). In the parietal ROIs, the ERP distribution was also asymmetrical with the contralateral hemisphere more positive. Additionally, a difference in ERP distribution between the left and right hand was found in the left-handed group, but not in the right-handed group (**Fig. 5B**). Three-way mixed ANOVA revealed a significant Hemisphere effect (F(1,20) = 5.708, p = 0.027, η_p_ ^2^ = 0.222), as well as a Hand x Handedness interaction (F(1,20) = 7.494, p = 0.013, η_p_ ^2^ = 0.273). Post hoc analysis showed left-handed participants had more positive parietal ERP when using the left hand (p = 0.041). Lastly, we performed the same ERP analysis by excluding the two participants with -33 and -7 EHI scores, and the results remained the same (see **Supplementary Materials**).

**Figure 5.**
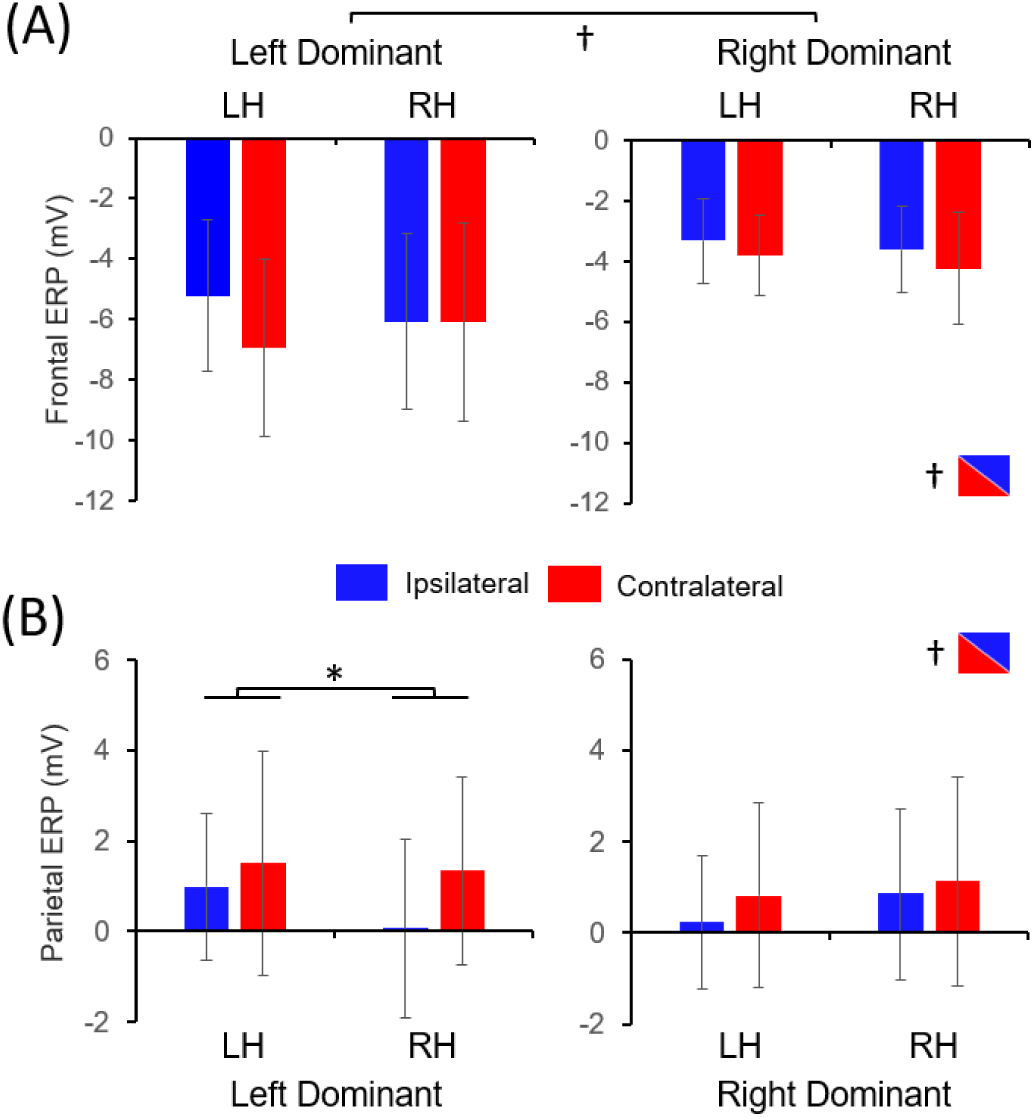
Comparisons of spatial ERP distribution (Mean ± S.D.) during N150 in (**A**) Frontal ROIs and (**B**) Parietal ROIs. LH and RH represent the left hand and the right hand as the acting hand. Daggers denote significant main effects (p < 0.05). The asterisk denotes a significant difference in post-hoc comparisons (p < 0.05).

### 3.4 Right-handed but not left-handed group showed hemispheric differences during P390

In the frontal ROIs, the left-handed group showed a symmetrical ERP distribution (**Fig. 4**). In contrast, ERP distributions were observed to be lateralized within the right-handed group. Specifically, the contralateral hemisphere showed positive ERP and the ipsilateral hemisphere showed negative ERP when the non-dominant left hand was used. However, when the dominant right hand was used, a positive ipsilateral ERP and a negative contralateral ERP were found. A 3-way mixed ANOVA revealed a 3-way *Handedness* x *Hand* x *Hemisphere* interaction (F(1,20) = 4.457, p = 0.048, η_p_ ^2^ = 0.182). Post hoc comparisons confirmed a significant difference between two hemispheres in both left-hand conditions (p = 0.017) and right-hand conditions (p = 0.027) for right-handed participants (**Fig. 6A**). No hemispheric difference was found for left-handed participants. In the parietal ROIs, it was found that the contralateral hemisphere ERP was generally more positive than the ipsilateral hemisphere ERP (**Fig. 4** and **Fig. 6B**). This was confirmed by a significant effect of *Hemisphere* in the 3-way mixed ANOVA (F(1,20) = 6.767, p = 0.017, η_p_ ^2^ = 0.253). Lastly, we performed the same ERP analysis by excluding the two participants with -33 and -7 EHI scores, and the results remained the same (see **Supplementary Materials**). Overall, these results suggest that during the P390 period, right-handers but not left-handers had a consistent rightward lateralization in the cortical activities, regardless of the hand performing the task.

**Figure 6.**
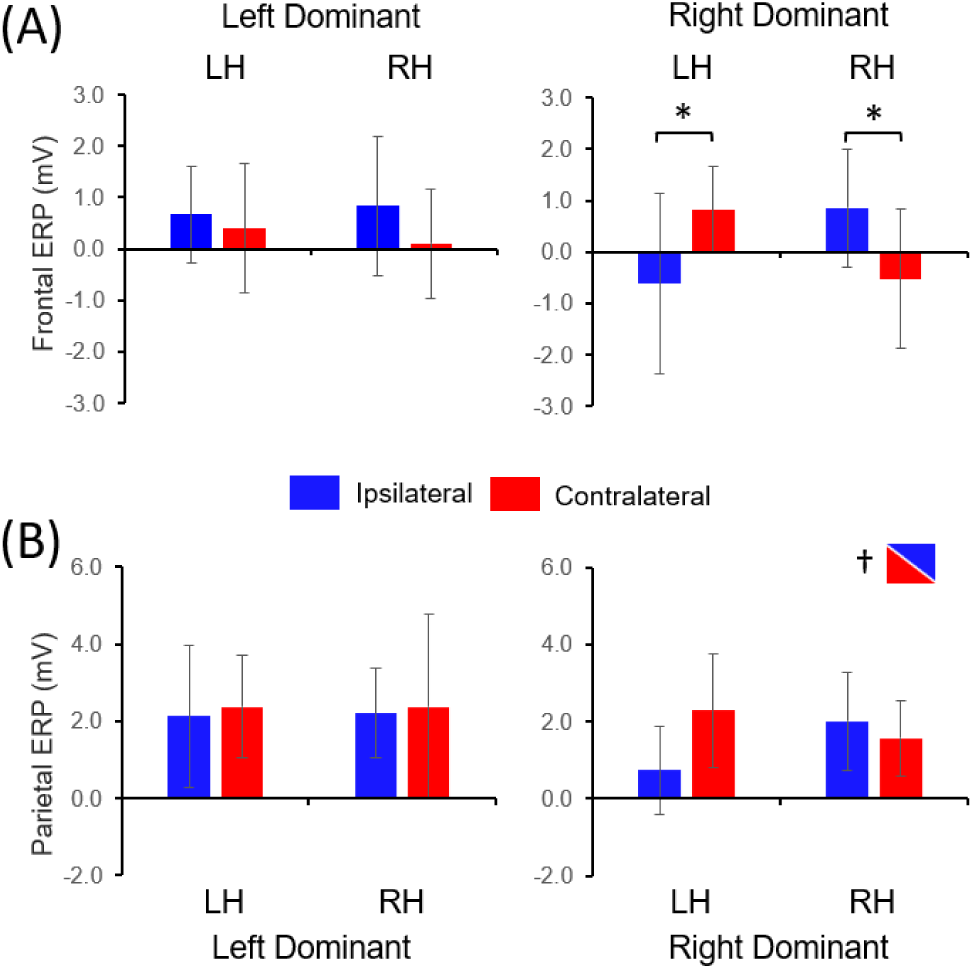
Comparisons of spatial ERP distribution (Mean ± S.D.) during P390 in (**A**) Frontal ROIs and (**B**) Parietal ROIs. LH and RH represent the left hand and right hand as the performing hand. Daggers denote significant main effects (p < 0.05). Asterisks denote significant differences in post-hoc comparisons (p < 0.05).

## 4. DISCUSSION

The present study examined both behavioral and neural responses after an unpredictable mechanical perturbation was applied in a simple reaching task. We did not find differences in movement kinematics between handedness groups or hand used for any conditions (**Fig. 2**). At first glance, this result seems surprising since there has been rich evidence that showed dominant and non-dominant arms exhibit asymmetric sensorimotor behavior across a wide range of tasks (Goble and Brown, 2008). For example, after unexpected inertial load changes, the non-dominant arm had greater final position accuracy than the dominant arm in right-handers (Bagesteiro and Sainburg, 2003), suggesting an advantage of the non-dominant arm in error correction driven by sensory feedback. However, the behavioral asymmetries between the two limbs were not always observed. It has been shown that two limbs exhibited similar behavior after perturbation during a posture maintenance task (Maurus et al., 2021). Moreover, the learning rates in force field perturbation schemes were not different between the right and left hands, even though the underlying adaptation mechanisms may be different (Schabowsky et al., 2007). Therefore, it is likely that specific task requirements are needed to discriminate the sensorimotor ability between two hands. For this study, participants were presented with a large Target sphere to move the cursor into, which does not require precise control of limb position causing similar movement kinematics. Despite the lack of behavioral differences, we observed significant differences in neural activities that are associated with both handedness and hand used. We discuss these important findings in relation to their functional relevance below.

### 4.1 Handedness effect on the cortical activity at 150 ms post mechanical perturbation

The onset of the three components of the responses to mechanical perturbation has been characterized by EMG (Kourtis et al., 2008; Pruszynski et al., 2008, 2011, 2014). The short latency component, which is a mono-synaptic spinal reflex, starts at 20-50 ms post perturbation. The long-latency component and the voluntary component start at 50-100 ms and 120+ ms post perturbation. Previous EEG studies using mechanical perturbation on the arms only focused on the early response period (0-100 ms post perturbation), which is characterized by an evoked potential located in the contralateral primary motor cortex with a peak at around 54-58 ms after perturbation onset (Mackinnon et al., 2000; Kourtis et al., 2008). This cortical response has been associated with the generation of the long-latency response, which was supported by intracortical recordings in non-human primate studies (Omrani et al., 2014; Pruszynski et al., 2014). We observed a similar cortical activity pattern during this early response period, an ERP component with strong frontal negativity and parietal positivity in the contralateral hemisphere at around 60 ms post-perturbation (**Fig. 3B** and **Fig. 4**). Additionally, we found no differences between the two handedness groups. However, the ERP after the initial 100 ms after perturbation onset has not been systematically investigated. Our results revealed that the ERP of left-handers exhibited greater negativity in the frontal area than the ERP of right-handers at around 150 ms post perturbation (**Fig. 5**). The topographical distribution of this N150 component suggests frontocentral cortical generators, with a small contralateral-ipsilateral asymmetry (**Fig. 3C)**. Moreover, given the timeline of the corrective response revealed by our behavioral data and previous EMG studies, this N150 component occurred after the long-latency response component, and it is likely associated with the process of voluntary control.

There have been studies showing the magnitude of frontal ERP can differ between left-handers and right-handers in some tasks, such as the P300 feature in visual or auditory oddball tasks (Alexander and Polich, 1997; Hoffman and Polich, 1999), and the P50 feature in a sentence completion task (Beratis et al., 2009). Kourtis and Vingerhoets (2016) demonstrated that right-handers exhibited a more negative frontocentral ERP peak at ∼ 125 ms and 250 ms after the presentation of a photo of an object that may be more suitable to grasp by one of the hands. The authors argued that the handedness effect on these frontal ERPs may be related to the working memory operation or attentional process, but the exact cortical generators that caused these handedness-related effects were not quantified. However, there are substantial differences between these studies and our study. First, these experimental tasks did not involve a perturbed environment in which online corrections were necessary. Second, the sensory modalities used in these tasks are mostly visual and auditory, whereas somatosensory information related to the mechanical perturbation is also important in our study. Third, the timings of the frontal ERP negative peaks that exhibited handedness effects in these tasks were different from that of our study (150 ms after the stimulus, i.e., perturbation). Therefore, we believe that the neural mechanism underlying our N150 component is unlikely to be the same as those that were speculated in the aforementioned studies.

Interestingly, the spatiotemporal pattern of our N150 component is very similar to a prominent ERP feature, commonly referred to as N1, that occurs after a mechanical perturbation during postural balance control tasks (Bolton, 2015). N1 plays an important role in the corrective response in the post-perturbation balance control. It has been shown that the magnitude of N1 scaled with the size of the perturbation (Mochizuki et al., 2010). Additionally, it has been shown that N1 is modulated by the predictability of the perturbation timing (Mochizuki et al., 2008), cognitive load (Little and Woollacott, 2015), and balance capability (Payne and Ting, 2020). These results indicated that N1 may be functionally relevant in the detection of postural instability. Source localization suggested that N1 is likely to originate from the supplementary motor area (SMA) with contributions from the anterior cingulate cortex (ACC) during postural tasks (Marlin et al., 2014; Mierau et al., 2015). Furthermore, a recent study showed that the amplitude of N1 after postural perturbation correlates with the amplitude of error-related negativity (ERN) during Flanker tasks (Payne et al., 2023), suggesting a shared neural network that combines SMA and ACC (Bonini et al., 2014) for action monitoring and error processing in a broad range of tasks. Based on these findings and the similarity in the experimental paradigms, we speculate that the functional role of N150 component in the present upper-limb perturbation paradigm is associated with the processes of error monitoring and action re-programming within the SMA/ACC area. Importantly, our results indicate a handedness-related difference in the neural activity of these processes. However, it remains to be determined whether such a handedness effect is caused by differences in morphology or neural dynamics between the two handedness groups. Future studies are also needed to examine the handedness effect on the N150 component across a broader range of motor tasks where online voluntary corrections are needed to correct errors induced by external perturbations.

### 4.2 Handedness effect on the cortical activity after 390 ms post mechanical perturbation

In reaching tasks, an abrupt visual shift of the target, i.e., target jump, before or at movement onset can elicit voluntary corrections with an onset time between 100 and 200 ms after the visual shift (Desmurget and Grafton, 2000; Veerman et al., 2008; Gritsenko et al., 2009). This timing is similar to the onset of the voluntary correction found in our study. By examining the spatiotemporal distribution of ERP in a target jump task, it was demonstrated that a positive ERP deflection peak over the parietal region occurs between 300 ms and 400 ms after the target jump in reaching performed by the dominant right hands of right-handers (Krigolson et al., 2008; Dipietro et al., 2013). These results are consistent with non-human primate intracortical recordings that showed significantly increased neuron spike rate in the parietal cortex during the same time window after a target jump in reaching (Archambault et al., 2011). It was theorized that this parietal activity may be associated with the update of the internal model of the environment after receiving new information estimated in the parietal circuits (Dipietro et al., 2013; Krigolson et al., 2015). Such an updated internal model can be important in predicting the consequence of actions in an environment altered by perturbations.

Despite the similarity in the timing between the ERP in the target jump paradigm and our P390 component, these studies did not discuss the symmetry of the ERP spatial distribution, making direct comparisons difficult. Additionally, it is unclear the extent to which they share a common functional role and neural mechanisms with our results, considering the differences in the source of online sensory feedback signals between visual and mechanical perturbations. A fMRI study has shown that cortical activation corresponding to target error (from visual target jump) and execution error (from mechanical curl field perturbation) during reaching may involve partially different networks in the parietal region, with the former extending more into the posterior areas (Diedrichsen et al., 2005).

Another (not mutually exclusive) explanation for the functional role of our P390 component, which lasted through the end of the trial, is the online control of end-point stabilization. Although our experimental paradigm did not have high precision requirements, the participants were instructed to stop within the target area at the end of each trial. Therefore, the last 100 ms of a trial, i.e., 400-500 ms post perturbation onset, likely involved decelerating and stabilizing the hand after the initial corrective actions in an altered environment (i.e., more or less force from the robot than before perturbation onset). This is supported by the relatively small changes in position as shown in **Fig. 2**. Therefore, it is plausible that the P390 component was related to a frontoparietal network that stabilizes the hand through online feedback control by integrating somatosensory and visual information (Suminski et al., 2007; Scott et al., 2015).

Interestingly, it has been proposed that different aspects of motor control may be associated with distinct neural lateralization patterns. The Dynamic Dominance Model has theorized that, in right-handers, the right hemisphere facilitates rapid online corrections, and the left hemisphere specializes in predictive feedforward control (Sainburg, 2014). This model aligns with a more general framework of neural lateralization where the left hemisphere is specialized for well-established patterns of behavior in familiar environments and the right hemisphere is specialized for responding to unforeseen events in uncertain environments (MacNeilage et al., 2009). Extensive behavioral studies revealed that the right arm is often better at producing more straight and efficient trajectories, whereas the left arm is better at stabilizing the hand as it approaches a target (Bagesteiro and Sainburg, 2002; Duff and Sainburg, 2007; Yadav and Sainburg, 2014). Given that a hemisphere contralateral to a given arm exerts a greater influence on that arm’s performance, these studies largely support the Dynamic Dominance Model. Moreover, it has been found that stroke survivors with left hemisphere damage had deficits in controlling the initial trajectory of fast motion, whereas patients with right hemisphere damage had deficits in maintaining end-point accuracy (Schaefer et al., 2007, 2009, 2012; Mani et al., 2013). These clinical data suggest that the lateralization of different movement control mechanisms is likely to occur in the same way for both arms in right-handers, i.e., hemispheric specialization, rather than in a mirrored fashion. Our result is consistent with the Dynamic Dominance Model of motor lateralization. Specifically, right-handers in the present study showed a rightward directional bias of the P390 ERP component for both the right hand and left hand (**Fig. 3D** and **Fig. 6**). This result indicates that the contributions of the frontoparietal networks in the right and the left hemispheres were consistently different for both hands in right-handers during the ending phase of voluntary corrections.

Studies examining the lateralization of motor control mechanisms in left-handers remain scattered. It was initially speculated that left-handers may have a lateralization pattern that mirrors the right-handers (Wang and Sainburg, 2006), i.e., the nondominant left hemisphere is specialized for feedback control of end-point position. However, this is only partially supported by behavioral data which also suggests that the lateralization of motor control may be less asymmetrical in left-handers than in right-handers (Przybyla et al., 2012; McGrath and Kantak, 2016). The Dynamic Dominance Model speculates that the behavioral symmetry of left-handers may be caused by bilateral hemisphere recruitment (Sainburg, 2016), which is in broad agreement with the handedness-related differences found in the spatial distribution of neural activities across various motor tasks that primarily rely on predictive control (Solodkin et al., 2001; Verstynen et al., 2005; Pool et al., 2014; Schmitz et al., 2019). In the present study, we extended previous research by showing that the symmetry of neural activities in left-handers was also different from that of right-handers in a task that primarily relies on feedback control.

Specifically, the P390 component showed a less lateralized pattern in left-handers than in right-handers (**Fig. 3D** and **Fig. 6**). This result suggests that the control of voluntary corrections in left-handers may involve a broader network distributed across two hemispheres, instead of relying on hemispheric specialization. Our finding is in general agreement with previously reported structural differences found through diffusion tractography, which revealed an effect of handedness on the asymmetry of frontoparietal tracts involved in integrating somatosensory and visual information but not on the asymmetry of corticospinal projection tracts (Howells et al., 2018).

### 4.3 Limitations

The present study has several limitations. First, our sample size (n = 22) is relatively small, which was only powered to detect large differences in ERP (see a-priori power analysis in **Supplementary Materials**). It is possible that we may not find subtle differences between left- and right-handed groups. Second, we did not use a pre-registration approach to constrain participant selection. This resulted in a higher heterogeneity of the left-handed group than the right-handed group. Moreover, handedness was treated as a binary variable rather than a continuous one, despite the heterogeneity within each handedness group. It is known that motor performance and the corresponding neural activity may be partly dependent on the consistency (or degree) of hand preference (Bernard et al., 2011; Davidson and Tremblay, 2013; Kourtis et al., 2014; Kourtis and Vingerhoets, 2016). Therefore, our results may be influenced by the difference in the handedness consistency between the two handedness groups. Future studies are needed to address the role of handedness consistency (i.e., correlation analysis with EHI scores), or the degree of hand preferences (i.e., strong versus weak), in the voluntary response to mechanical perturbation. Third, we only used EHI to measure hand preferences but did not quantify hand performance in other simple motor tasks. Past studies have shown that hand performance and hand preference are only moderately correlated (Triggs et al., 2000; Brown et al., 2004; Chatagny et al., 2013). It was argued that limb preferences may result from the interaction between lateralization of motor control processes and task constraints (Sainburg, 2014), therefore only using hand preferences in the present study may be oversimplifying the underlying neural lateralization. Future studies should consider classifying participants with a multimodal approach that depends on both hand preference and hand performance metrics.

Lastly, we were not able to perform source localization with our data because the relatively small number of electrodes and the lack of individual electrode location data prevented accurate estimation of dipole locations. With only data at the channel level, we can only infer the underlying cortical processes based on existing research and the fact that a source near a recording electrode tends to produce stronger signals than sources further away from the electrode. While we acknowledge that our interpretation should be taken with caution, our results provide important insights into the differences in the sensorimotor control of voluntary correction after mechanical perturbation between left- and right-hand dominant individuals.

## 5. CONCLUSION

In summary, for the first time, the present study demonstrated that frontoparietal activities of left-handers and right-handers differ in response to mechanical perturbations of upper limbs. In left-handers, we observed a greater negative deflection of ERP in the frontal region at 150 ms after perturbation onset. Moreover, we found that left-handers showed a more symmetrical ERP spatial distribution than right-handers after 390 ms after perturbation onset.

These ERP differences suggest that the neural circuits underlying the voluntary corrective sensorimotor control of the right and left hands, especially for error monitoring and online movement stabilization, may differ between left- and right-handers.

## Glossary

EEG: Electroencephalography ERP: Event-related potential

EHI: Edinburgh handedness inventory TANOVA: topographic analysis of variance ROI: Regions of interest

SMA: Supplementary motor area ACC: Anterior cingulate cortex

## Author contributions

Conceptualization and methodology: K.H., K.K, Q.F.; Data collection: K.H. and K.K; Data analysis: K.H., K.K, Q.F.; Data visualization: K.H. and Q.F.; Manuscript writing, review, and editing: K.H. and Q.F.

## Acknowledgments

The authors would like to thank Joseph Dranetz and Maya Patel for their assistance with data collection.

