## Supplementary material for "Cortical neural activity during responses to mechanical perturbation: Effects of hand preference and hand used"

**Manuscript title:**

**Supplementary Materials**

### 1. Power analysis

During the design phase of the study, we performed a priori power analysis to determine the sample size using G*Power software (Faul et al., 2007). The effect size was estimated from the literature that used similar designs. For example, Kourtis& Vingerhoets (2016) analyzed ERP in both left- and right-handed individuals who responded to photos of graspable objects. They used a three-way ANOVA with Handedness and Handedness consistency as two between factors, and hemisphere as a within factor. They reported partial eta squared values ranging from 0.155 to 0.402 (Cohen's f ranging from 0.4 to 0.8). Similar effect sizes were also reported elsewhere (Schmitz et al., 2019). Therefore, we used $f = 0.5$, $\alpha = 0.05$, and $\beta = 0.8$ to determine the sample size for our three-way mixed ANOVA. It was estimated that 22 participants were needed for the main effect of the between factor (i.e., Handedness), and fewer were needed for the within factors and interactions.

### 2. EHI results of all participants.

The Edinburgh Handedness Inventory (EHI) asks the hand preferences in the following tasks: writing, drawing, throwing, scissors, toothbrush, knife, spoon, broom (upper hand), striking match, open a lid. One or two points are assigned to one hand if that hand is preferred or strongly preferred, respectively. One point is assigned to each hand if there is no preference. The EHI score is determined following the formula [(R-L)/(R + L)]*100, with R denoting the sum of right-hand points and L denoting the sum of left-hand points, leading to a Laterality Quotient (L.Q.) scores between -100 and 100.

| Right-handed | | Left-handed | |
|---|---|---|---|
| Sub ID | L.Q. | Sub ID | L.Q. |
| 2 | 88 | 1 | -80 |
| 3 | 100 | 5 | -62 |
| 4 | 100 | 15 | -7 |
| 6 | 90 | 16 | -33 |
| 7 | 88 | 17 | -57 |
| 12 | 65 | 18 | -60 |
| 13 | 70 | 21 | -100 |
| 14 | 90 | 22 | -75 |
| 19 | 100 | 23 | -46 |
| 20 | 100 | 24 | -40 |
| | | 25 | -73 |
| | | 26 | -87 |

The table above summarizes all participants' L.Q. scores. A cut-off of 0 can be used to differentiate left- and right-handed participants (Oldfield, 1971). However, L.Q. scores can represent various degrees of handedness consistency, and past studies have used other cut-off scores (Edlin et al., 2015). For example, an L.Q. between -40 and 40 can be considered ambidextrous (Giovagnoli & Parisi, 2024).

**3. Results without ambidextrous participants.**

The following results were obtained with the same methods described in the main manuscript, but two participants with L.Q. scores between -40 and 0 were excluded. Note that these results were similar to the results presented in the main manuscript.

3.1 The kinematic responses were similar across handedness groups and acting hands

We did not find significant main effects of Handedness and Hand or their interactions on the perturbation-related or target-related kinematic response differences.

3.2 Spatial distributions of ERP were different across handedness groups.

TANOVA revealed that a significant main effect of Handedness was found between 110 ms and 160 ms, as well as between 390 ms and 440 ms post perturbation onset. Significant main effects of Hand were found for all 10 ms time windows after 40 ms post perturbation onset. A significant interaction was only found between 320 ms and 340 ms after perturbation onset.

3.3 The left-handed group had more negative frontal ERP during N150.

In the frontal ROIs, we observed that the left-handed group showed a greater negative peak in the ERP during N150. Moreover, the N150 was more negative in the contralateral hemisphere, suggesting an asymmetrical distribution of ERP with respect to the midline. This was confirmed by a three-way mixed ANOVA showing a significant effect of Handedness ($F(1,18) = 7.258$, $p = 0.015$, $\eta_p^2 = 0.287$) and Hemisphere ($F(1,18) = 17.328$, $p < 0.001$, $\eta_p^2 = 0.490$). In the parietal ROIs, the ERP distribution was also asymmetrical with the contralateral hemisphere more positive. Additionally, a difference in ERP distribution between the left and right hand was found in the left-handed group, but not in the right-handed. Three-way mixed ANOVA revealed a significant Hemisphere effect ($F(1,18) = 6.975$, $p = 0.017$, $\eta_p^2 = 0.279$), as well as a Hand x Handedness interaction ($F(1,18) = 7.220$, $p = 0.015$, $\eta_p^2 = 0.286$). Post hoc analysis showed left-handed subjects had more positive parietal ERP when using the left hand ($p = 0.045$).

3.4 The right-handed group showed hemispheric differences during P390.

In the frontal ROIs, the left-handed group showed a symmetrical ERP distribution. In contrast, ERP distributions were observed to be lateralized within the right-handed group. Specifically, the contralateral hemisphere was positive, and the ipsilateral hemisphere showed negative ERP when the non-dominant left hand was used. However, when the dominant right hand was used, a positive ipsilateral ERP and a negative contralateral ERP were found. A 3-way mixed ANOVA revealed a 3-way *Handedness* x *Hand* x *Hemisphere* interaction ($F(1,18) = 5.234$, $p = 0.034$, $\eta_p^2 = 0.225$). Post hoc comparisons confirmed a significant difference between the two hemispheres in both left-hand conditions ($p = 0.007$) and right-hand conditions (p =

0.007) for right-handed subjects. No hemispheric difference was found for left-handed subjects. In the parietal ROIs, it was found that the contralateral hemisphere ERP was generally more positive than the ipsilateral hemisphere ERP. This was confirmed by a significant effect of *Hemisphere* in the 3-way mixed ANOVA ($F(1,18) = 7.299$, $p = 0.015$, $\eta_p^2 = 0.289$). Overall, these results suggest that during the P390 period, right-handers but not left-handers had a consistent right-ward lateralization in the cortical activities, regardless of the hand performing the task.

**4. No perturbation conditions**

The main focus of this study was the voluntary corrective responses after mechanical perturbation was applied. Nevertheless, we had two No Perturbation conditions (UN and DN) in the randomized trial sequence to maximize the unpredictability of the Perturbed conditions. They were not analyzed in the main manuscript, but we present the results of these two conditions here.

4.1 Movement kinematics. The data was analyzed using an approach similar to the analysis used in the main manuscript. We did not find a significant effect of Handedness or Hand. Overall, it was estimated that the movements toward the targets started at approximately 260 ms (Fig. S1).

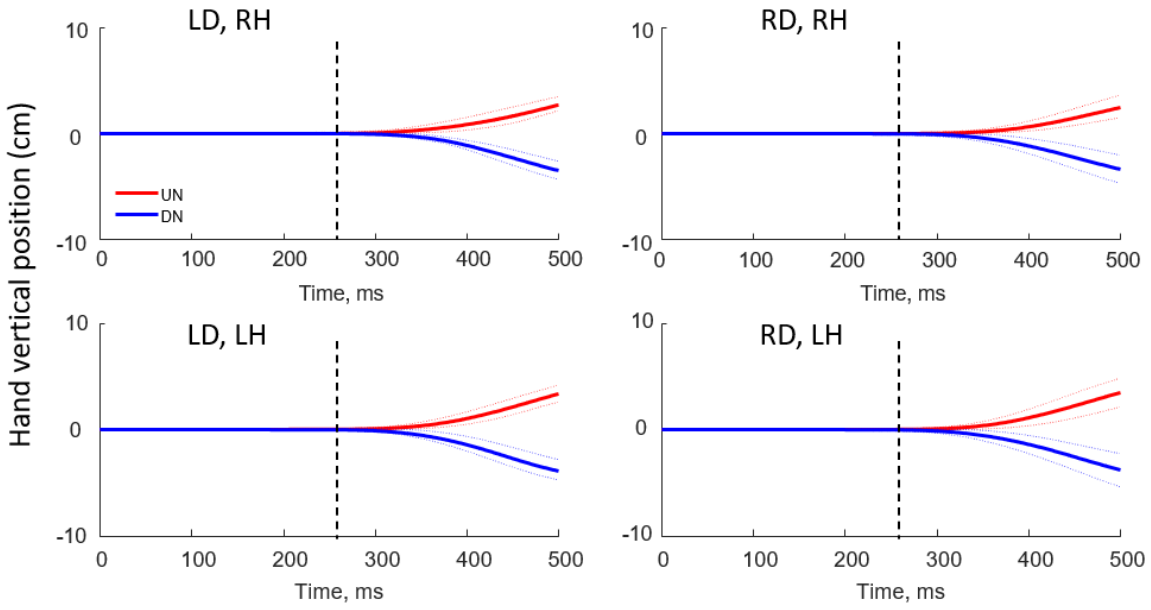


**Figure S1**. Kinematics of the no perturbation conditions UN and DN (mean ± S.D.). LD and RD denote Left- and Right-handed groups, respectively. RH and LR denote the right hand and left hand as the acting hand in the task. The vertical dashed line represents the estimated onset of the voluntary movement.

4.2 TANOVA results.

We did not find an effect of Handedness in the overall distribution of ERP in the period of 0 – 500 ms post perturbation. There were a few points in the period where we found a main effect of Hand (Fig. S2), which was most likely caused by the motor planning and execution processes primarily occurring in the hemisphere contralateral to the acting hand.

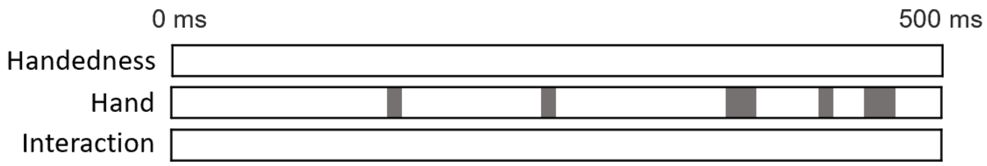


**Figure S2**. Each horizontal bar represents results from TANOVA analysis for the 0 – 500 ms period post perturbation onset. Dark-shaded areas indicate significant effects ($p < 0.05$).

4.3 ROI results.

We did not perform statistical analysis with the ROI data because no effect of Handedness was found in the TANOVA. Nevertheless, we present the ERP signals from the frontal and parietal ROI in Figure S3 (Left-handed group) and S4 (Right-handed group).

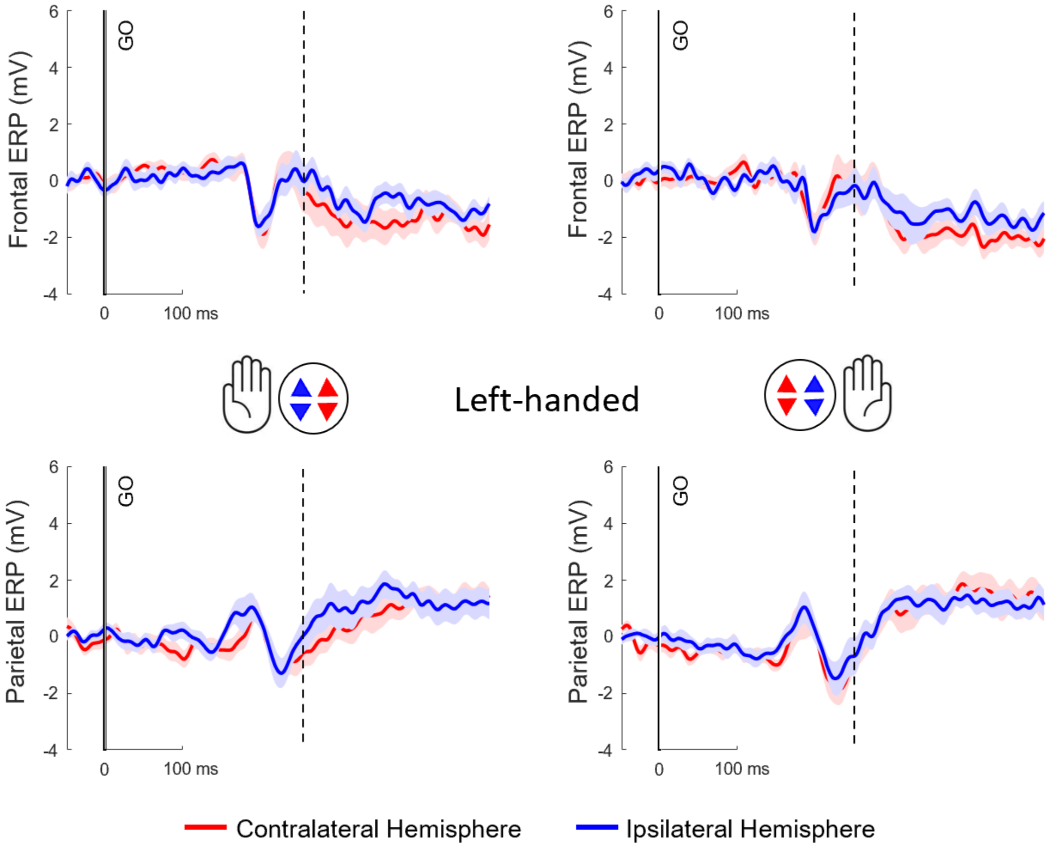


**Figure S3**. ERP traces from frontal ROIs (top row) and parietal ROIs (bottom row) of Left-handed individuals. Red lines and shaded areas (Mean ± S.D.) represent the contralateral hemisphere, blue lines and shaded areas represent the ipsilateral hemisphere. The vertical dashed line represents the estimated movement onset.

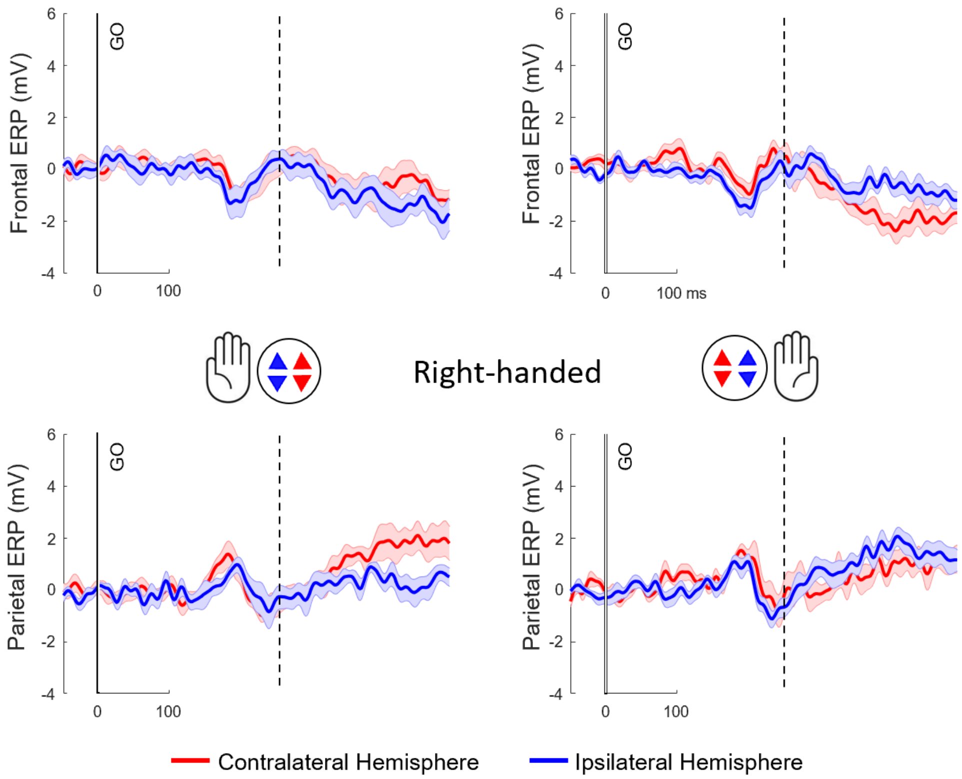


**Figure S4**. ERP traces from frontal ROIs (top row) and parietal ROIs (bottom row) of Right-handed individuals. Red lines and shaded areas (Mean ± S.D.) represent the contralateral hemisphere, blue lines and shaded areas represent the ipsilateral hemisphere. The vertical dashed line represents the estimated movement onset.
